# Dynamic Laser Speckle Imaging meets Machine Learning to enable Rapid Antibacterial Susceptibility Testing (DyRAST)

**DOI:** 10.1101/2020.02.04.926071

**Authors:** Keren Zhou, Chen Zhou, Anjali Sapre, Jared Henry Pavlock, Ashley Weaver, Ritvik Muralidharan, Joshua Noble, Jasna Kovac, Zhiwen Liu, Aida Ebrahimi

## Abstract

Rapid antibacterial susceptibility testing (RAST) methods which measure change of a bacterial phenotype in response to a given treatment are of significant importance in healthcare, as they can assist care-givers in timely administration of the right treatment. Various RAST techniques have been reported for tracking bacterial phenotypes, such as size, shape, motion, and metabolic activity. However, they still require bulky and expensive instruments (which hinders their application in resource-limited environments) and/or utilize labeling reagents (which can interfere with antibiotics and add to cost). Furthermore, the existing ultra-rapid methods do not address possible adaptation of gradual adaptation of bacteria to antibiotics, which can lead to false interpretation of resistance when using ultra-rapid methods. In this work, we present a RAST approach leveraging machine learning analysis of time-resolved dynamic laser speckle imaging (DLSI) results to accurately predict the minimum inhibitory concentration (MIC) of a model strain of *Escherichia coli* in 60 minutes, compared to 6 hours using the currently FDA-approved phenotype-based RAST technique. To demonstrate the DLSI performance, we studied the effect of a *β*-lactam ampicillin and an aminoglycoside gentamicin on *Escherichia coli* strain K-12. DLSI captures change of bacterial motion/division in response to treatment. The machine learning algorithm was trained and validated using the overnight results of gold standard, broth microdilution method. Empowered by machine learning, DyRAST can predict MIC with high accuracy comparable to gold standard methods through a voting strategy.

## Introduction

Antimicrobial resistance (AMR) is among the most serious health threats of all times. Antimicrobially resistant pathogens cause estimated 2.8 million infections and 35,000 deaths per year in the United States.^1^ The fatality rate due to antimicrobially resistant infections is expected to reach 10 million per year by 2050 if the transmission of AMR is not effectively mitigated.^2^ Misuse and overuse of broad-spectrum antibiotics due to lack of reliable and accurate rapid antibacterial susceptibility testing (RAST) are contributing to growing occurrence of antimicrobially resistant infections. Gold standard AST techniques, including disk diffusion and broth microdilution (BMD), take over 18 hours to complete, which limits their utility in cases of severe sepsis.^3,4^ Development of accurate RAST that produce results concordant with gold standard methods is therefore critically needed to speed up AST to support data-based antibiotic prescribing and improve patient treatment outcomes.

A series of molecular and phenotypic RAST technologies have recently been developed. Genotypic RASTs rely on the detection of target nucleic acid sequences from pathogens to determine and predict the antibiotic resistance.^5^ Amplification of target DNA using polymerase chain reaction (PCR),^6^ isothermal recombinase polymerase amplification (RPA),^7^ or loop-mediated isothermal amplification (LAMP),^8,9^ combined with fluorescence detection^10^ can provide the results within an hour in a single assay. Due to diverse genetic resistance determinants that do not have sufficient correlation with resistance phenotypes, culture-independent AST testing cannot be broadly applied to all pathogens without extensive prior knowledge of their biology and underlying genetics. Furthermore, molecular methods still fail to detect resistance in cases where novel resistance mechanisms have not yet been characterized.^11–13^

Compared to molecular methods, phenotype-based ASTs in many cases provide information that is more relevant to clinical outcomes. Among different phenotypic methods, Accelerate Pheno™ (Accelerate Diagnostics, Inc., Tucson, AZ, USA) is currently the only FDA-approved RAST technique that measures morphokinetic cellular changes that occur due to exposure to antibiotics. This method combines fluorescence in situ hybridization (FISH) and automated microscopic imaging to obtain AST results within 6-7 hours.^14^ A method reported by Schoepp *et al*. has further improved the time to results by determining antimicrobial susceptibility of *E. coli* directly from clinical urine samples in 30 min.^8^ Their technique, coupled with digital LAMP, measures microbial growth based on quantification of nucleic acids using quantitative PCR. One of the limitations of this ultra-fast technique is assuming that slowed or halted DNA replication after 15 min antibiotic exposure indicates susceptibility of a bacterial population to a given antibiotic. However, the bacterial population response within the first 15 min is typically highly variable among assay replicates, which increases an error in prediction of susceptibility,^15^ especially when borderline MIC concentrations of antibiotics are tested. In addition to these two approaches, other RAST methods measure changes in bacterial viability in response to antibiotics by probing other phenotypic features, including shape,^16^ motion,^17^ size/mass,^18,19^ or respiration.^20^

Among phenotypic RAST, methods which rely on monitoring bacterial motion have been extensively explored. Change of bacterial viability in response to antibiotics can also be measured by probing other phenotypic features, including shape,^16^ motion,^17^ size/mass,^18,19^ or respiration.^20,21^ Among them, AST methods which rely on monitoring bacterial motion are particularly attractive. Johnson *et al*. used the phase noise of resonant crystal to prove that the motion of *E. coli* was attenuated after treated with polymyxin B or ampicillin.^22^ This work makes the bacterial motion as a promising characterization for RAST. Yu *et al*. reported a phenotypic AST method that utilized deep learning to analyze the freely moving *E. coli* cells in urine and can determine the minimum inhibitory concentration (MIC) in 30 mins.^16^ However, these methods have poor sensitivity and/or require labeling^23^ or indicative markers such as redox-active chemicals, which can interfere with cellular physiology.^20^ In addition, these methods still need advanced imaging/analysis setups, such as high resolution optical microscopy, atomic force microcopy, or complex optical setups,^16,24^ which limits their broad application in clinical diagnostics.

Compared to optical methods that visually assess changes in shape, length, and/or motion of bacterial cells, measuring light scattering off of bacterial populations is a simpler method that eliminates the need for advanced optical setup. One of such methods is laser scattering, which has been utilized in developing RAST. For example, BacterioScan system (BacterioScan Inc.) measures optical density (OD) of a liquid sample as well as the low-angle laser scattered intensity which enables measuring significantly lower OD levels compared to traditional ratiometric transmittance measurement.^25^ However, this method is based solely on OD, which does not provide for substantial improvement of the RAST turnaround time (~ 6 hrs).

In order to further reduce the detection time, we utilized dynamic laser speckle imaging to quantify bacterial micromotion and correlate the inactiveness in micromotion with the inhibitory effects of antibiotics. The methods is based on the phenomenon of laser light scattering in turbid media, such as biological tissues.^26^ The interference of the scattered light produces a speckle pattern that can convey characteristics of the scatterers and the nature of their behavior over time. The differentiation between dead and alive bacterial cells is achieved based on the dynamic analyses of the laser speckle images. When the light is scattered from a static turbid medium, a constant laser speckle pattern is generated due to the fixed phase profile of the scattered light from a static medium.^27–30^ On the contrary, if particles in the medium exhibit intrinsic movement, the scattering pattern will changes over time, generating a dynamic speckle pattern.^29^ A temporal series of speckle images hence produces patterns that contain information about particles’ kinetic behavior and can be used as a means to understand the effect of environmental triggers, including antibiotics, on their motion. For example, Murialdo *et al*. used dynamic laser speckle imaging (DLSI) to detect different degrees of motility and chemotaxis in bacteria swarming plates.^28^ Ramirez-Miquet *et al*. proposed a technique combining speckle imagining with a digital image information technology (DIT) to track multiplying *E. coli* and *S. aureus* cells deposited on agar plates at high concentration of 1.5×10^9^ cell/mL and 10^9^ cell/mL, respectively.^29^ They compared the mean viability of each pathogen and showed that speckle imagining method has a potential to detect biological activity in 15 mins. Despite these efforts, to the best of our knowledge, there is no report of an application of time-resolved dynamic laser speckle imaging for antibacterial susceptibility testing in liquid samples. Moreover, previously reported assays are not sensitive enough for analysis of cells at approximately 5× 10^5^ cell/mL, a concentration required by the gold standard protocols.

Here, we show that analysis of the time-resolved dynamic laser speckle images using artificial neural network (ANN) was able to identify the MIC of ampicillin and gentamicin for *E. coli* K-12 in only 60 minutes with high accuracy comparable with gold standard methods using a voting strategy for the ANN predictions. The predictions were validated using gold standard broth microdilution. Further work based on this proof of concept is needed to evaluate the method’s performance on a broader panel of microorganisms and antibiotics. DyRAST is a rapid, label-free phenotypic AST technique that utilizes simple optical instrumentation. Dynamic speckle imaging eliminates the need for advanced microscopy systems, which significantly simplifies the RAST. Moreover, the developed method has a potential for adjustment to operate using consumer-level components, such as smartphone camera and off-the-shelf laser diodes.

## Materials and Methods

### Bacterial culture

*Escherichia coli* (*E. coli*) strain K-12 was used as a model bacterial strain in all experiment. The culture was stored as a frozen stock at −80 °C and resuscitated every 14 days to maintain a fresh inoculum. The culture was resuscitated by streaking onto a Brain Heart Infusion (BHI) agar and then incubated at 37 °C 20+/-2 h. A single colony from a BHI agar was inoculated into the 10 mL BHI broth and incubated at 37 °C, shaking at 210 rpm, for 20+/-2 h. The overnight culture was diluted in Mueller Hinton Broth (MHB) based on the OD600 to obtain 5 × 10^5^ CFU/mL using a BioPhotometer D30 (Eppendorf, Hauppauge, NY). An OD600 of 1 was assumed to be equal to 8 × 10^8^ CFU/mL.

### Preparation of the antibiotics

Ampicillin (Sigma Aldrich, CAS# 7177-48-2) and gentamicin (Sigma Aldrich, CAS# 1405-41-0) were prepared by dissolving an antibiotic powder in sterilized MilliQ ultrapure water to achieve 5 mg/mL and 10 mg/mL stock solutions, respectively. All the stock solutions were frozen in 0.1 mL aliquots and stored at −20 °C until use antimicrobial susceptibility testing.

### Antimicrobial susceptibility testing using broth microdilution

The culture in concentration 5 × 10^5^ CFU/mL was used for broth microdilution, speckle imaging, and broth macrodilution (i.e., time-kill kinetics). For broth microdilution, the standard methods recommended by the Clinical and Laboratory Standards Institute (CLSI) guideline M100-S22^31^ were followed to determine the MIC of ampicillin (AMP) and gentamicin (GEN) for *E. coli* strain K-12. Briefly, 50 μL of MHB were aliquoted in wells of 96-well microtiter plates (Greiner bio-one). Antibiotics were added to the wells of the first row and sequentially two-fold diluted. Fifty microliters of culture prepared as described above were added to each well, and incubated for 16-20 hrs. Negative and positive controls were included in each test plate and each test was carried out in a triplicate in at least two independent experiments. MIC was determined by visually inspecting wells for turbidity resulting from a culture growth.

### Antimicrobial susceptibility testing using broth macrodilution

To determine the time-kill kinetics using broth macrodilution, culture was sampled and viable cells were quantified at times 0, 1, 2, 3, 5, 8, 16, and 24 h by spiral plating (easySpiral, Interscience) 10-fold dilutions onto MHB agar. Inoculated plates were incubated at 37 °C for 16-20 h and counted to determine the CFU/mL. A negative control that was not treated by an antibiotic was included in each experiment. After an initial 1 hour incubation of bacterial cells in MHB (to reach early exponential growth phase), antibiotics with different concentrations are added to separate cuvettes. Control culture without antibiotics was also tested. The cuvettes were subsequently placed into the custom-built holder in the optical setup to capture dynamic speckle patterns. All experiments are done in triplicates.

### Dynamic laser speckle imaging (DLSI) setup

The optical measurement system consists of a laser source, a lens, a cuvette holder and a camera (Fig. 1). Distances between the components of the system are listed in the Table S1, in the Supplementary Information, SI. We have optimized the setup based on the Mie scattering model described in SI, Section 2. A helium-neon laser (wavelength: 632.8 nm, power: 0.8 mW, HNLS008L, Thorlabs Inc., USA) was used as an illumination source. The laser beam was slightly expanded by using a concave lens (focal length= −25.0 mm) before illuminating a cuvette (Fisherbrand, CAS#14-955-129) containing 3 mL of bacterial suspension. The resultant speckle pattern was captured by a CMOS camera (Zyla, ANDOR), that was controlled by a computer.

**Fig. 1.**
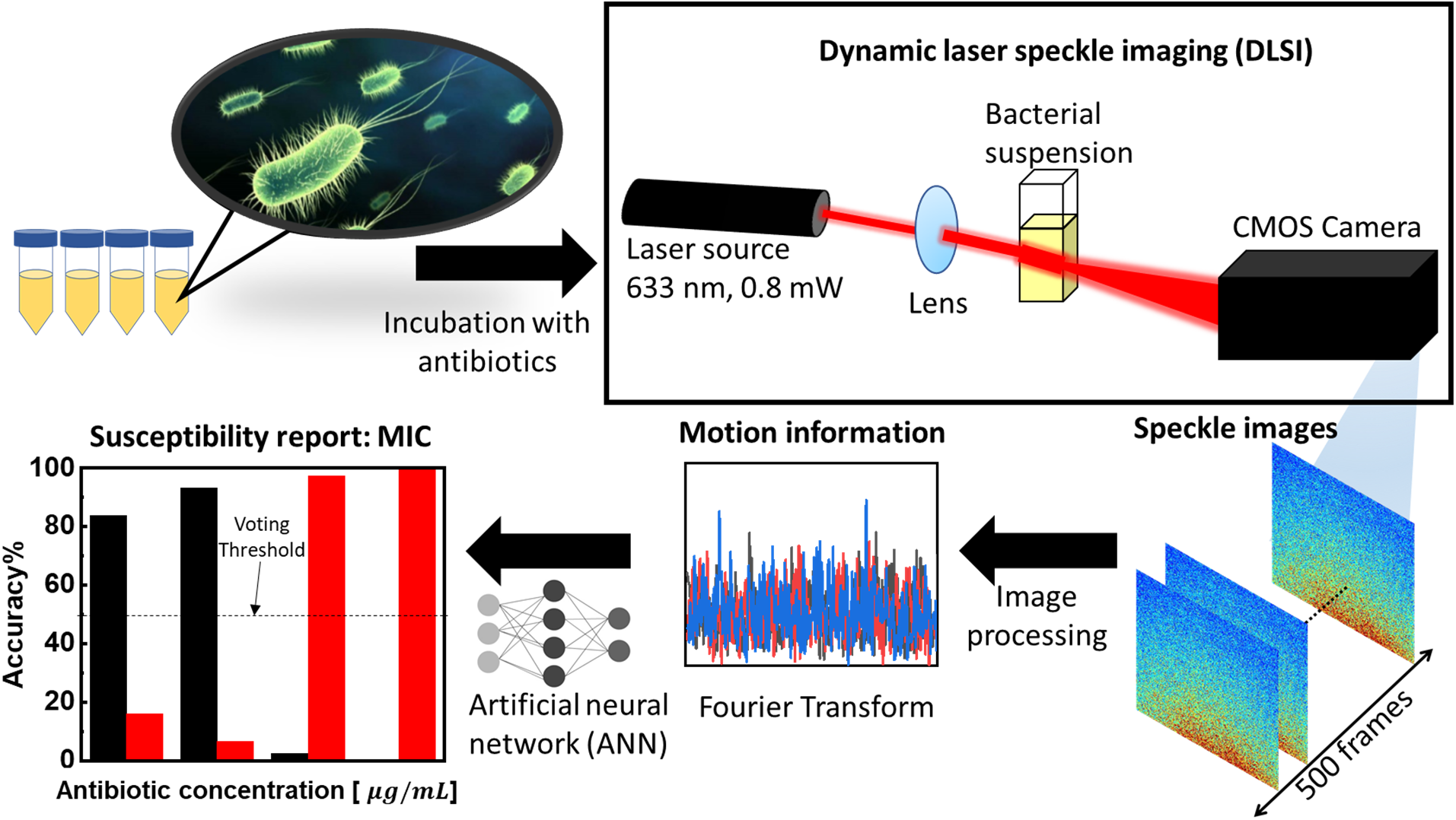
Schematics of the setup of dynamic speckle imaging for rapid AST (DyRAST). After expanded by the lens, the laser beam is scattered by the bacterial culture in the cuvette containing different concentration of antibiotic (*ρ_Antb_*). At each time point (*t_i_*), the dynamic speckle patterns were recorded, for total time of 10 sec, 500 frames. After pre-processing, Fourier analysis reveals the information about bacterial motion. The Fourier results are then fed into an artificial neural network (ANN). Finally, the trained ANN model can accurately predict the MIC in 60 min.

### Antibacterial susceptibility testing using DLSI

For DLSI-based AST, the culture adjusted to 5 × 10^5^ CFU/mL was further incubated at 37 °C, 210 rpm for 1 h to reach the logarithmic growth phase before an antibiotic was added to the culture in concentration equal to 0×MIC (control), 0.5×MIC, 1×MIC, and 2×MIC. Three mL of each culture was collected at each time point, *t_i_* (*i* = 0,1,…, 4) at 0, 30 min, 60 min, 90 min, and 120 min, respectively, and transferred into the cuvette used for DLSI in the imaging system described above. At each *t_i_*, a dynamic speckle patterns series (16 bits, 1000 × 2000 pixels, 50 frames per second) were collected. The exposure time for each frame was 1 msec (millisecond) and the full capture period was 10 sec at each time point, *t_i_*. All experiments were conducted in independent triplicates.

### Image pre-processing and machine learning model

The raw images (1000 × 2000) were first resized (100 × 200) using the nearest neighbor method to reduce the computational cost. The camera has a pixel size of 6.5 μm, leading to a 6.5 *mm* × 13 *mm* detection area. Fourier Transform (FT) was performed along the time axis of the measured data cube. Only the spectral intensity was used in our analysis since a spectral intensity distribution with significant high frequency content generally corresponds to more active motion, whilst the spectral phase is not directly linked to motion activeness and presents a challenge in quantitative interpretation. This procedure transforms an original 100 × 200 × 500 data cube (two-dimensional space and one-dimensional time) into a new 100 × 200 × 249 data cube consisting of 249 positive frequency sampling points at each pixel position. The DC term was first normalized to 1 for each individual pixel spectrum, and was subsequently removed. Moreover, the negative frequencies provide identical information as their positive counterparts. Essentially, each measurement captures 20,000 spectra and each of the spectra contains 249 frequency features. This large amount of data provides a basis for machine learning based analytics.

Artificial Neural Network (ANN) with a hidden layer containing 300 neurons was constructed for classification prediction. Input layer had 249 neurons, representing the frequency components, up to 25 Hz. The output was defined as 2 classes, “Inhibited” and “Non-Inhibited”, referring to bacterial susceptibility or resistance due to exposure to antibiotic. The stochastic gradient descent (SGD) method was used to minimize the binary cross-entropy loss function. Binary classification for the “Inhibited” and “Non-Inhibited” bacteria groups was determined with the input from frequency domain for each pixel. All the image pre-processing and machine learning computations were performed with a desktop computer (Intel Core i9-9900k, 32GB RAM).

## Results and Discussion

To identify the setup parameters affecting the DLSI sensitivity, we adjusted and studied the effect of the distance between the laser, lens, cuvette, and camera (see Fig. S1 and Table S1 in the Supplementary Information, SI). We investigated three configurations: Setting #1-3. The important parameter in the setup is the angle between the axis of the camera and the laser beam (*θ*) which affects the speckle intensity, as described by the Mie scattering model (detailed discussion in SI, Section 2). The setup with Setting #3, which collects the speckle patterns at 11°-22° scattering angle, provides high sensitivity for laser speckle imaging (consistent with the Mie scattering model) in the current study, while also prevents the direct incidence of the laser beam on the camera. The results reported here are based on this configuration.

Fig. 2 shows typical speckle patterns of *E. coli* culture captured at *t_i_* = 0, 30, 60, 90, and 120 min with different concentrations of ampicillin and gentamicin (*ρ_AMP_* and *ρ_GEN_*), with respective MIC values of 4 *μg/mL* and 2 *μg/mL*. The speckle patterns for the control experiment (without antibiotics) are shown in the first row. In the absence of antibiotics, bacteria cells are healthy and can multiply. Their multiplication and division translate into an increase of the total number of scatterers, and hence increase of the intensity of the speckle patterns over time. With sub-inhibitory MIC concentration (e.g. 0.5×MIC value shown in the second and fourth rows, for AMP and GEN, respectively), initially there is an increase of the speckle intensity, followed by a slight decrease at 120 min. When cells are exposed to MIC (the third and fifth rows, for ampicillin and gentamicin, respectively), there is no significant change of the speckle intensity over time, which indicates inhibition of bacterial division.

**Fig. 2.**
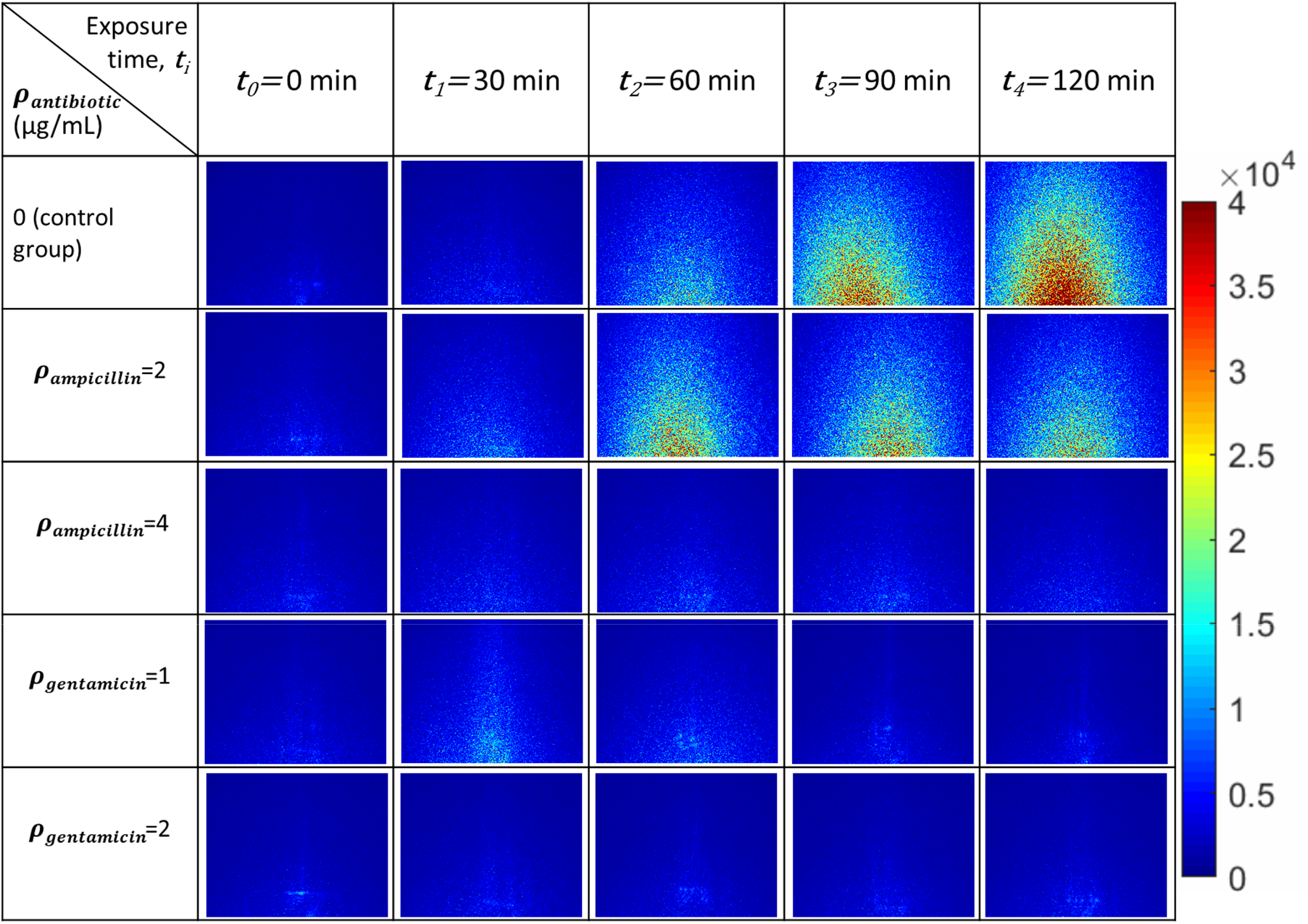
Raw representative speckle images of *E. coli* K-12 exposed to ampicillin (AMP) and gentamicin (GEN) at different concentrations (*ρ*). MIC of ampicillin and gentamicin was 4 *μg/mL* and 2 *μg/mL*, respectively. The images were collected over a total period of 2 hrs with 30 min intervals at time points, *t_i_* (*i* = 0,1,…, 5). The images were collected using the optimized setup (Setting #3).

The OD data and the average intensity values obtained from raw speckle patterns (examples shown in Fig. 2) are shown in Fig. 3a-b and Fig. 3c-d for ampicillin and gentamicin, respectively. The average intensity for each raw speckle image was obtained on each pixelated image matrix collected at *t_i_’*s (hence, obtaining a data cube with dimension of Spatial: 1000 × 2000 × #Time frames: 500). Average intensity obtained using DLSI captures time-kinetics of *E. coli* growth more sensitively than conventional OD measurements (with detection limit indicated by dashed lines in Fig. 3). While the OD values for different antibiotic concentrations are below the detection limit of the OD reading even after 120 min, the average speckle intensity values are distinguishable in 90 min. This demonstrates the advantage of DLSI (even before applying machine learning) compared to the existing techniques that measure optical absorption.

**Fig. 3.**
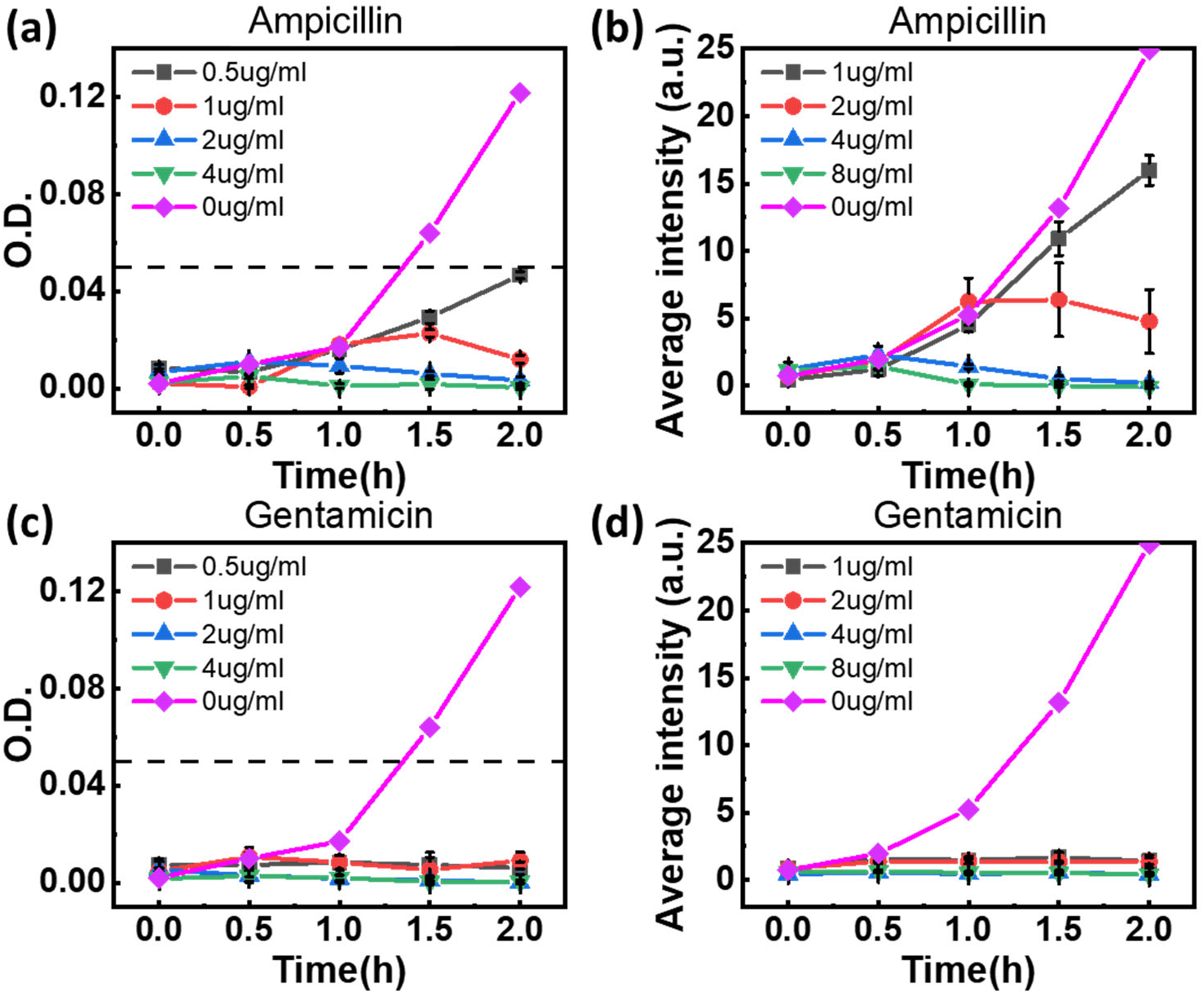
(a) The OD_600_ values vs. treatment time and (b) the average intensity results obtained from raw dynamic laser speckle images of *E. coli* K-12 with ampicillin with *ρ_AMP_* = 0,1,2,4, and 8 *μg/mL*. (c) and (d): The corresponding results for gentamicin with *ρ_GNT_* = 0,0.5,1,2 *and* 4 *μg/mL*. The dashed lines in part (a) and (c) indicate the detection limit of the commercial OD measurement system. The MIC for ampicillin and gentamicin were 4 *μg/mL* and 2 *μg/mL*, respectively.

However, simple intensity measurement is “blind” to the overnight (> 16 h) turbidity/colony count results (Fig. 3a/Fig. 5a and Fig. 3c/Fig. 6a for AMP and GEN, respectively), and hence prone to false positive; for example, determining ampicillin MIC to be 2 *μg/mL* (Fig. 3), while the correct MIC based on OV data is 4 *μg/mL*. The intensity measurement also neglects the time-resolved motility information of cells. Moreover, absolute intensity is sensitive to setup specification (such as laser power, exposure time, camera gain, etc.), and hence present challenge for calibration and expansion to a general case. In order to overcome this problem, we utilize machine learning and train artificial neural network (ANN) models using the OV gold-standard results to analyze the dynamic speckle patterns to determine MIC. Fig. 4 demonstrates our methodology for preprocessing the data and the machine learning algorithm. As mentioned earlier, we optimized the setup based on Mie scattering model to collect the speckle patterns at the angular range of 11°-22°. To this end, the transmitted laser beam is blocked to avoid saturation of the camera and ensure collection of the scattered light. As shown in Fig. 4a, Fourier Transform of individual pixels was obtained, with the DC term normalized to 1 (details provided in SI, Section 3) so that the analysis is independent on the absolute speckle intensity. For each time point of each group, we obtained 20,000 pixel-level spectra, each of which consists of 249 frequency features. We then built a separate ANN model for each time point, with 249 neurons as the input, 300 hidden units, and 2 output neurons for binary classification.

**Fig. 4.**
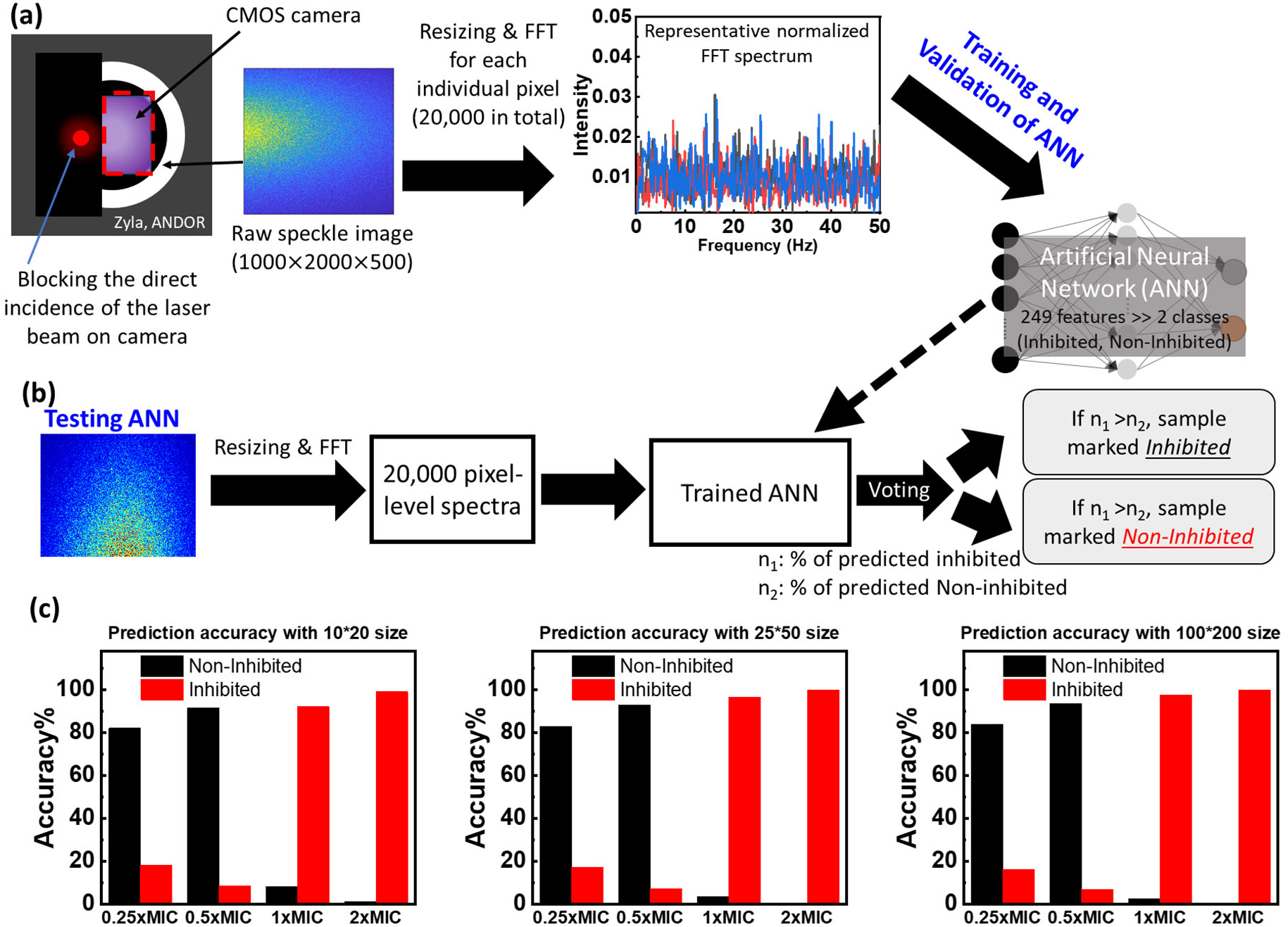
Data preprocessing and machine learning algorithm for prediction of the minimum inhibitory concentration (MIC) and whether a particular antibacterial treatment is inhibitory or not. (a) Raw speckle images were collected at each *t_i_*. The incident laser beam was blocked to ensure it does not directly hit the camera to avoid pixel saturation. The time series at each pixel was Fourier transformed with the DC term normalized to 1. Fourier results, i.e., the resultant features, were fed into the ANN model, with 249 neurons as the input, 300 hidden units, and 2 output neurons for binary classification. (b) The trained ANN model was tested using a separate set of data. Similarly, 20,000 pixel-level spectra were preprocessed and fed into the trained ANN. Through voting (threshold at 50%), we determined the susceptibility of the bacterial sample (“Inhibited” vs. “Non-Inhibited”). (c) Comparison of the ANN algorithm with different resizing factor. Prediction accuracy of ANN with 100×200, 25×50, or 10×20 resizing factor is almost identical (within 5% difference).

**Fig. 5.**
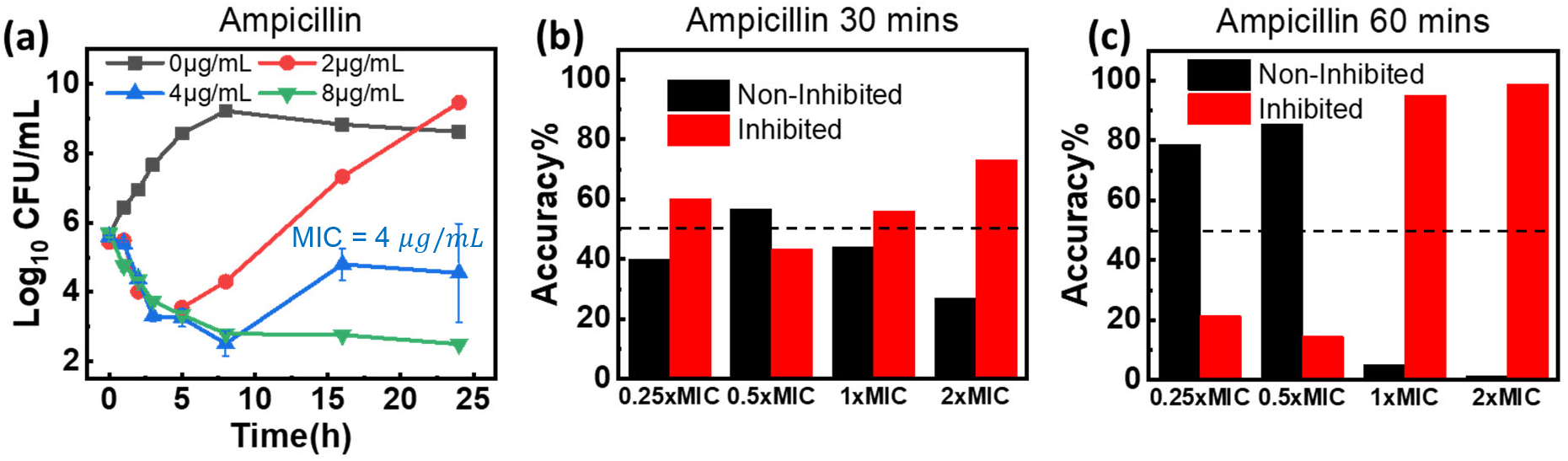
(a) The time-kill curves for *E. coli* K-12 with different concentrations of ampicillin (*ρ_AMP_*). The prediction accuracy of machine learning algorithm improves with time: (b) the results using DLSI data collected at *t_2_* =30 min, and (c) the results using DLSI data collected at *t_3_* =60 min. After 60 min, the method can identify MIC with high accuracy. The dashed lines indicate the voting threshold (50%) to predict AST results with high accuracy, i.e. “Inhibited” for 1×MIC and 2×MIC, while at the same time, “Non-Inhibited” for 0.25×MIC and 0.5×MIC. Ampicillin MIC was 4 *μg/mL*.

**Fig. 6.**
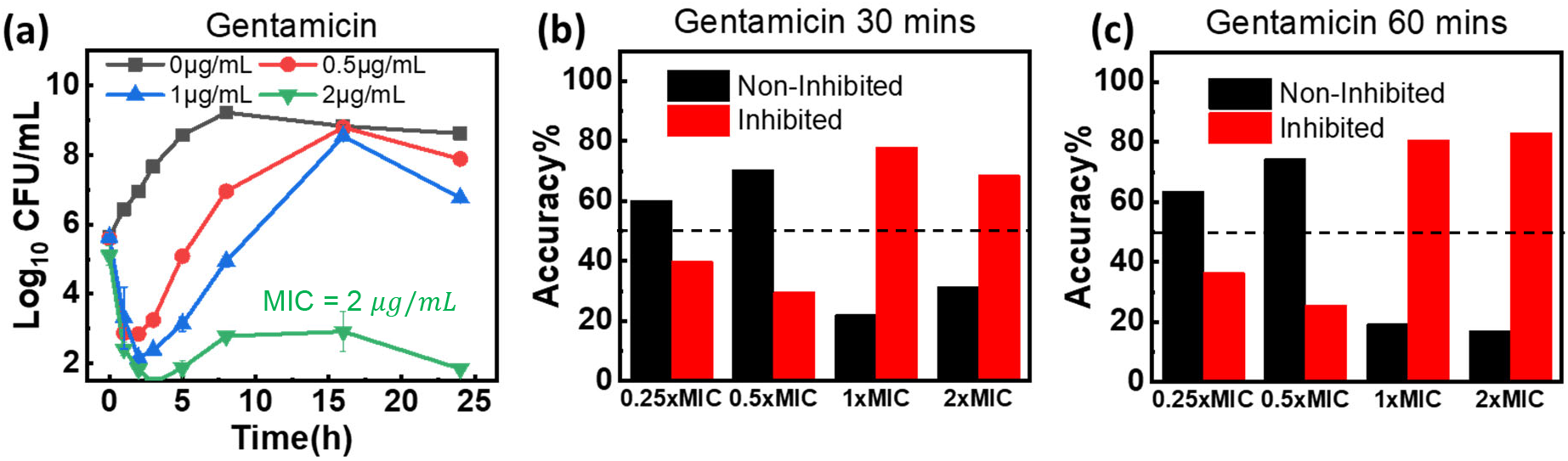
(a) The time-kill curves for *E. coli* K-12 with different concentrations of gentamicin (*ρgen*). The prediction accuracy of machine learning algorithm improves with time: (b) the results using DLSI data collected at *t_2_* =30 min, and (c) the results using DLSI data collected at *t_3_* =60 min. Interestingly, the method can predict MIC for gentamicin in 30 min (compared to 60 min for AMP). The dashed lines indicate the voting threshold (50%) to predict AST results with high accuracy, i.e. “Inhibited” for 1×MIC and 2×MIC, while at the same time, “Non-Inhibited” for 0.25×MIC and 0.5×MIC. Gentamicin MIC was 2 *μg/mL*.

We trained the ANN models with experimental data with *E. coli* K-12 exposed to antibiotics of different concentrations (0.25×, 0.5×, 1× and 2×MIC). We conducted experiments in 3 independent replicates with groups exposed to four concentrations for each antibiotic. The four concentrations yielded a total of 80,000 spectra for each time point. As shown schematically in Fig. 4b, we split the samples randomly to training, validation, and test groups with a ratio of 70%, 15%, and 15%, respectively. The validation was carried out to avoid model overfitting, and the test group was used to verify the predictive power of the trained neural networks. Two independent experiments where each experiment was completed with four concentrations of the antibiotic resulted in a combined data size of 160,000 samples at each time point, were used in building the learning models. We then used a third, independent experiment that consisted of 80,000 spectra at each time point for a final comprehensive test to evaluate the robustness and accuracy of trained models.

The overnight OD values were utilized to label the dataset. If the OD after overnight culture incubation was at the same level of the initial OD, the bacterial group was labeled as “Inhibited”. On the other hand, if the final OD was higher than the initial OD value (above the resolution limit of the equipment which is ~0.05), indicating that the antibiotic did not inhibit the growth of the bacteria, the group was labeled as “Non-Inhibited”. In our experiments, 0.25× and 0.5×MIC were labeled as “Non-Inhibited”, while 1× and 2×MIC were labeled as “Inhibited”. For each group at a specific time point, *t_i_*, since all the 20,000 pixel-level spectra have the same label, the result of the test should be interpreted as a voting process where the percentage of each prediction category indicates the likelihood for this group to be classified as the said category. For example, 30% predicted as “Inhibited” means that 30% out of the 20,000 spectra are predicted as “Inhibited”, while the remaining 70% is predicted to be “Non-Inhibited”. Therefore, this group is classified as “Non-Inhibited.” As shown in Fig. 4c, we also evaluated the effect of the resizing factor on performance (prediction percentage) of the machine learning analysis. The blue bar indicates the percentage of pixels that were predicted “Non-Inhibited”, while the orange bar indicates the percentage of “Inhibited”. With 100×200, 25×50, or 10×20 resizing factors, there was only 5% difference among the machine learning results, and they are all able to correctly predict MIC in 60 min through the voting strategy using a 50% threshold for decision-making. The ability to reduce the data size using a higher resizing factor indicates that computational cost can be reduced without significant performance degradation. It should be noted that the ML method does not rely on the absolute speckle intensity which could be difficult to calibrate, as the DC frequency component is normalized to 1. Detailed discussion on machine learning and confusion matrices (which allow visualization of the ANN performance) are provided in SI, Section 4.

The machine learning results for prediction of ampicillin and gentamicin susceptibility and MIC are shown in Fig. 5 and 6, respectively. Fig. 5a and Fig. 6a show the time-kill curves obtained using broth macrodilution method. The pixel-level prediction percentage obtained using machine learning are shown in Fig. 5b-c and Fig. 6b-c, for AMP and GEN, respectively. Fig. 5b suggests that 30 min is not long enough for accurate prediction of MIC. We define final classification result based on a voting strategy: the dashed lines in Fig. 5 and 6 indicate the voting threshold to predict antimicrobial susceptibility. We expect a classification “Inhibited” for 1×MIC and 2×MIC, while at the same time, we expect to a classification “Non-Inhibited” for 0.25×MIC and 0.5×MIC. For example, at 30 min, the pixel-level predicted percentage of classification into “Inhibited” or “Non-Inhibited” for 2×MIC of AMP is approximately 50%, making it difficult to decide which category the sample is more likely to fall into. While at 60 min, the correct category (“Inhibited”) is predicted in more than 80% of the votes. Interestingly, by comparing Fig. 5 and 6, it is observed that the method can predict MIC for gentamicin in 30 min (compared to 60 min for AMP). As a result, in order to confidently identify MIC in current experiments, a minimum of 60 min is required by DyRAST for accurate analysis. The pixel-level prediction percentage difference increases with time, as shown in SI, Fig. S8 and Fig. S9 for ampicillin and gentamicin, respectively.

It is worth noting that by counting viable cells using the broth macrodilution technique (a sensitive yet slow and laborious method), differentiation among 0.5×MIC, 1×MIC and 2×MIC is possible only after 5 hours for ampicillin and after 2 hours for gentamicin (see Fig. 5a and 6a). The only alternative rapid phenotype-based method available on the market and approved by the FDA, requires at least 6-hour culture incubation to confidently determine the antimicrobial susceptibility.^14^ DyRAST can accurately determine an MIC of ampicillin and gentamicin for *E. coli* K-12 within 1 hour of culturing an isolate. Table 1 compares the performance of DyRAST reported in this study with other phenotypic RASTs based on the minimum cell concentration, RAST time, sample condition, and setup complexity. We also compared the methods based on the complexity of the setup that is utilized.

**Table 1.** A comparison of this work with some of the recent phenotype-based RAST methods.

| Method | Cell concentration | Sample | AST time | Validation method | Setup complexity | Reference |
| --- | --- | --- | --- | --- | --- | --- |
| **Accelerate PhenoTest BC Kit** | $10^5$ cell/mL | Clinical specimens | 6h | Compared with disk diffusion result | High (require labeling) | <sup>14</sup> |
| **OCelloScope** | $2\times 10^7$ cell/mL | Veterinary samples | 3h | N/A | Low | <sup>32</sup> |
| **Drast** | $5 \times 10^5$ to $5 \times 10^7$ cell/mL | Bacteria add into fresh blood | 6h | Compared with BMD | High (Microscopy) | <sup>33</sup> |
| **SNDA-AST** | $5 \times 10^5$ cell/mL | Clinical isolates and urine samples | < 5.5h | N/A | High (Microscopy) | <sup>1</sup> |
| **Stress-based Microfluidic System** | 100-4000 cells in 12 nL | Four Gram-negative strains with MHB | < 1h | MIC Determined using VITEK<sup>®</sup> | High (Microscopy) | <sup>34</sup> |
| **Laser Ablation Electrospray Ionization Mass Spectrometry** | 1.5–2.5 cells per 1000 $\mu\text{m}^2$ | Disk | 21h | Compared with MIC database | High (Microscopy) | <sup>35</sup> |
| **Nanomotion sensor** | OD= 0.5 | Clinical *B. pertussis* | 3h | Compared with BMD | High (AFM<sup>b</sup> setup) | <sup>36</sup> |
| **Nanomotion-AST** | 100 cell/mL | Bacteria with human blood | < 3h | N/A | High (AFM setup) | <sup>37</sup> |
| **BacterioScan** | $10^5$ cell/mL | Bacteria with MHB<sup>a</sup> | 6h | N/A | Low | <sup>25,38</sup> |
| **DyRAST** | $5 \times 10^5$ cell/mL | *E. coli* K12<br>with MHB | 1h | Compared with<br>BMD | Low | **This work** |

<sup>a</sup>MHB: Mueller-Hinton Broth; <sup>b</sup>AFM: Atomic Force Microscopy

## Conclusion

This work presents a rapid, phenotype-based antibacterial susceptibility testing method capable of identifying MIC in 60 minutes. The method leverages machine learning analysis of time-resolved dynamic laser speckle imaging patterns to predict antimicrobial susceptibility and MIC in a rapid and reliable manner. The DLSI data was collected using a simple-to-use, low-cost optical setup, with no labeling or advanced imaging/optical setup required. To demonstrate the capabilities of the method (termed as DyRAST), we studied the effect of two antibiotics, ampicillin and gentamicin, which have different mechanisms of action. DLSI captures change of bacterial motion/division in response to antibiotic treatment. The DyRAST was validated against the gold standard AST methods using *E. coli* K-12 as a model microorganism. By adapting a voting strategy for analysis of the pixel-level prediction values obtained using artificial neural network models, the method predicted MIC with high accuracy comparable with gold standard methods in all tested conditions. The technique can be optimized for analysis of other (pathogenic) bacterial and fungal species and their response to antimicrobial treatment. Moreover, considering no complicated optical components are required for speckle imaging, a portable version of DyRAST can be potentially adapted by using consumer-level components, such as smartphone camera and laser diodes.

## ACKNOWLEDGMENTS

The authors would like to thank the Materials-Life Science Convergence Award supported by the Penn State’s Materials Research Institute, Huck Institute of Life Sciences, College of Engineering, and College of Medicine (Hershey). ZL and CZ acknowledge the support by Penn State MRSEC, Center for Nanoscale Science, under the award NSF DMR-1420620. JK was supported by the USDA National Institute of Food and Agriculture Hatch Appropriations under Project #PEN04646 and Accession #1015787.

