## Supplementary Information for "Dynamic Laser Speckle Imaging meets Machine Learning to enable Rapid Antibacterial Susceptibility Testing (DyRAST)"

### Section 1: Setup configuration

***Table S1.*** *The optical modules position parameters.*

| Setting | a | b | c | θ |
|---|---|---|---|---|
| **#1** | 9 cm | 6 cm | 8 cm | 20° |
| **#2** | 9 cm | 6 cm | 4 cm | 10° |
| **#3** | 9 cm | 5 cm | 6 cm | 15 ° |

*a: distance between the lasers and lens. b: distance between the lens and cuvette. c: distance between the cuvette and camera. θ: the angle between the camera optical axis and the laser beam.*

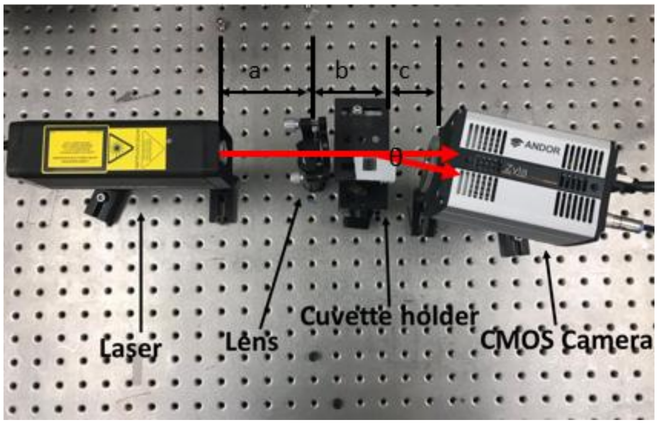


**Fig. S1.** The picture of the test setup with component position parameters.

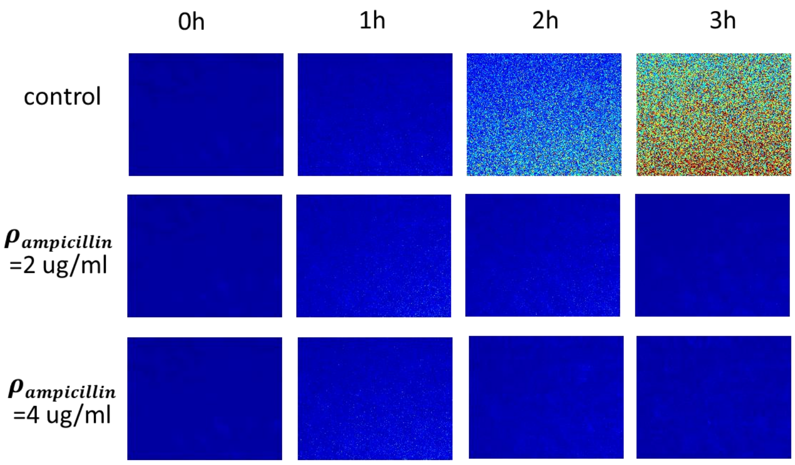


**Fig. S2.** Raw speckle images obtained using Setting#1. In this case, the scattered light is too weak (consistent with the Mie scattering model) for pattern recognition by machine learning.

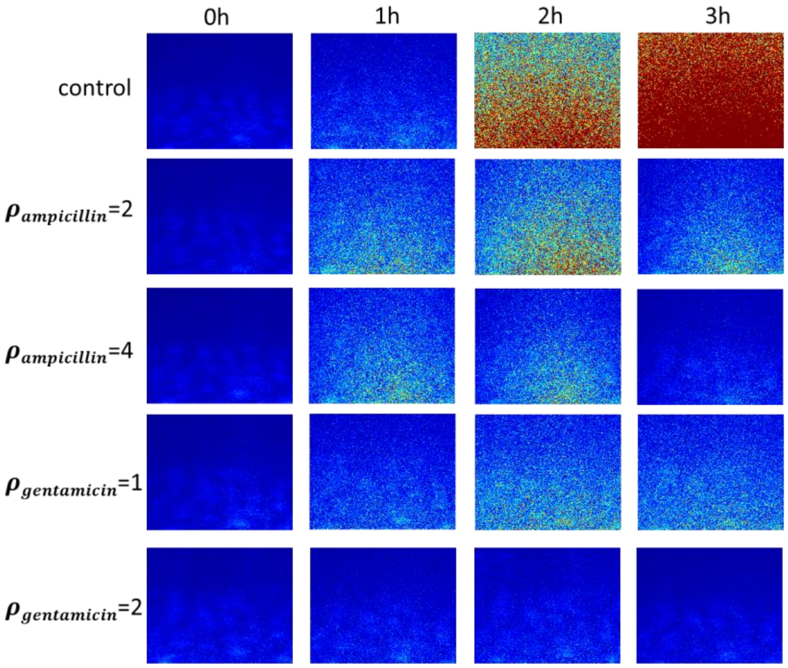


**Fig. S3.** Raw speckle images obtained using Setting#2. In this case, the signals can saturate due to high intensity of the scattered light.

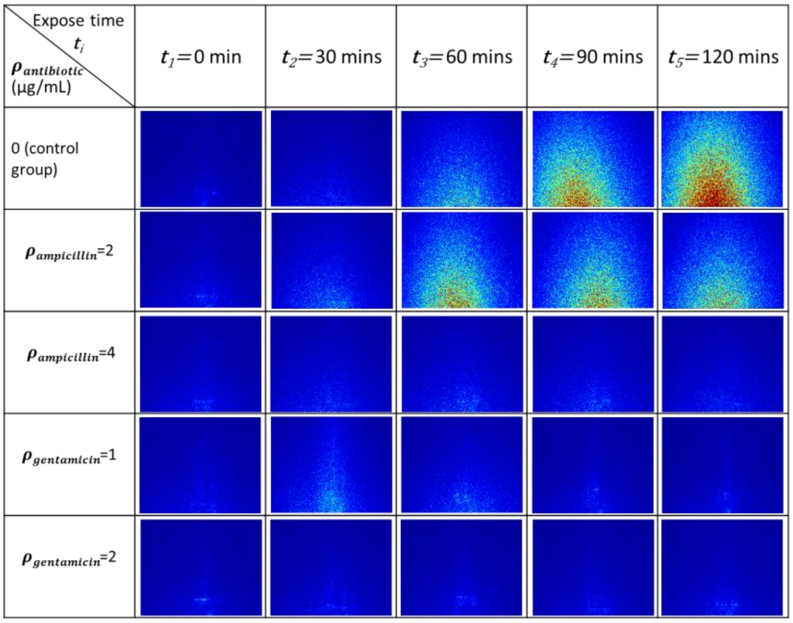


**Fig. S4.** Raw speckle images obtained using Setting#3 (gaining the best results based on ML analysis).

### Section 2: Mie scattering analysis

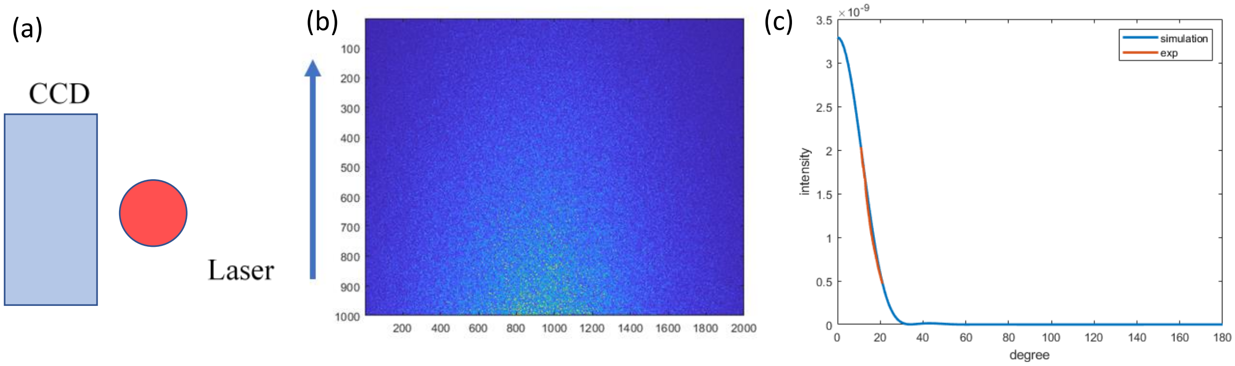


**Fig. S5.** (a) The location of captured window of CCD relative to the laser beam. A speckle pattern has 1,000 pixels in lateral direction (increasing scattering angle), and 2,000 in the vertical direction. (b) The representative speckle image. The blue arrow indicates the lateral direction. (c) The simulated result from Mie scattering model. Assume n(*E. coli.*) = 1.384, n(water) = 1.33, particle radius = 0.5 micro meter, the laser wavelength = 633 nm. The angular dependence of the intensity is shown. In our experiment, an angular range between 11 degree to 22 degree was measured, which is shown in the orange line plot.

### Section 3: Fourier Transform (FT) analysis

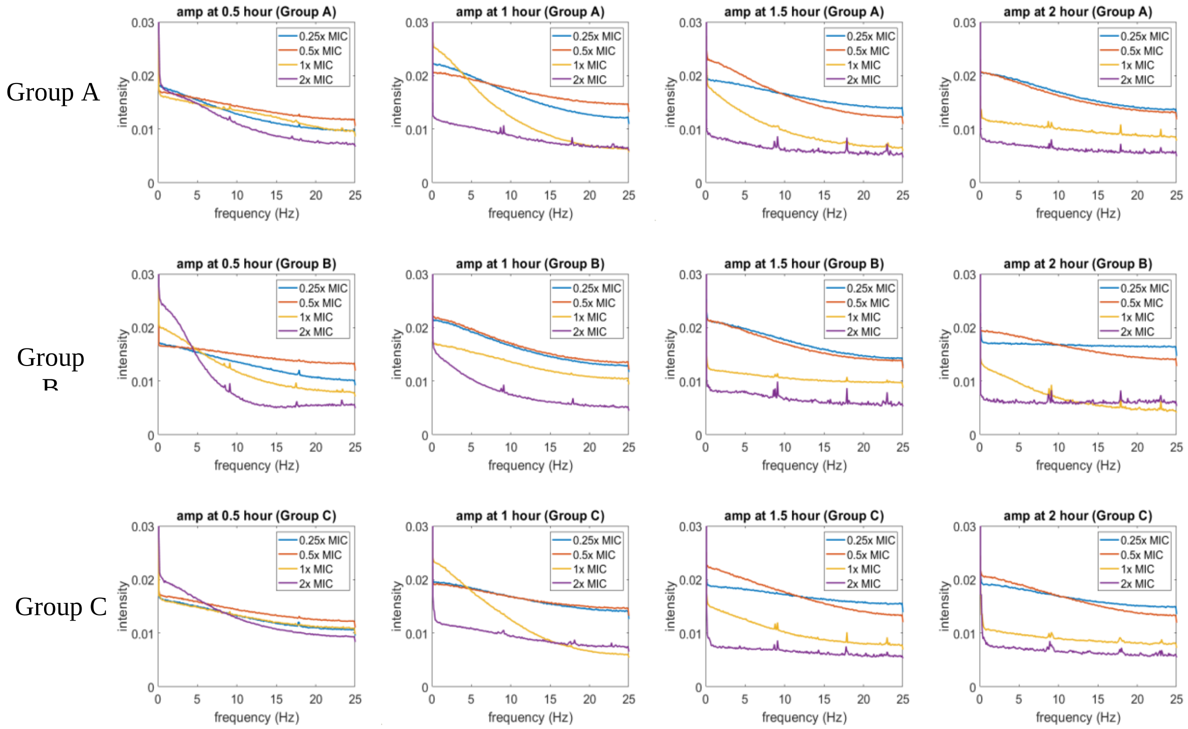


**Fig. S6.** The FT result of ampicillin group for three independent experiments at time points of 30m, 1h, 90 min, and 2h. The rows correspond to different experiments and the columns indicate different time points.

The DC frequency component is normalized to 1 for each individual pixel after the FT operations. For 100x200 resizing factor, we have 20,000 samples for each group. The average FT curve is calculated and plotted in Figure S6. In our experiments, we have up to 25 Hz bandwidth. After 1.5 hours, we notice that the 0.25x and 0.5x MIC groups, which are resistant groups, have more high frequency contributions compared to the 1x and 2x MIC, which are the susceptible groups. This indicates resistant groups have more active motion than the susceptible groups. However, we cannot clearly distinguish the difference at 0.5 hour, and 1 hour from the averaged curves. From the FT analysis, we conclude that it contains information about the mobility of the bacteria and provides features that machine learning algorithms can analyze and make predictions. Data obtained using the gentamicin group are processed with the same analysis methods as ampicillin. As shown in Figure S10, 0.25x and 0.5x MIC show higher frequency component. Similar to ampicillin results, the variance of individual pixel spectra is significant. We utilized machine learning to make pixel-level predictions and then apply a voting strategy for prediction of MIC and susceptibility.

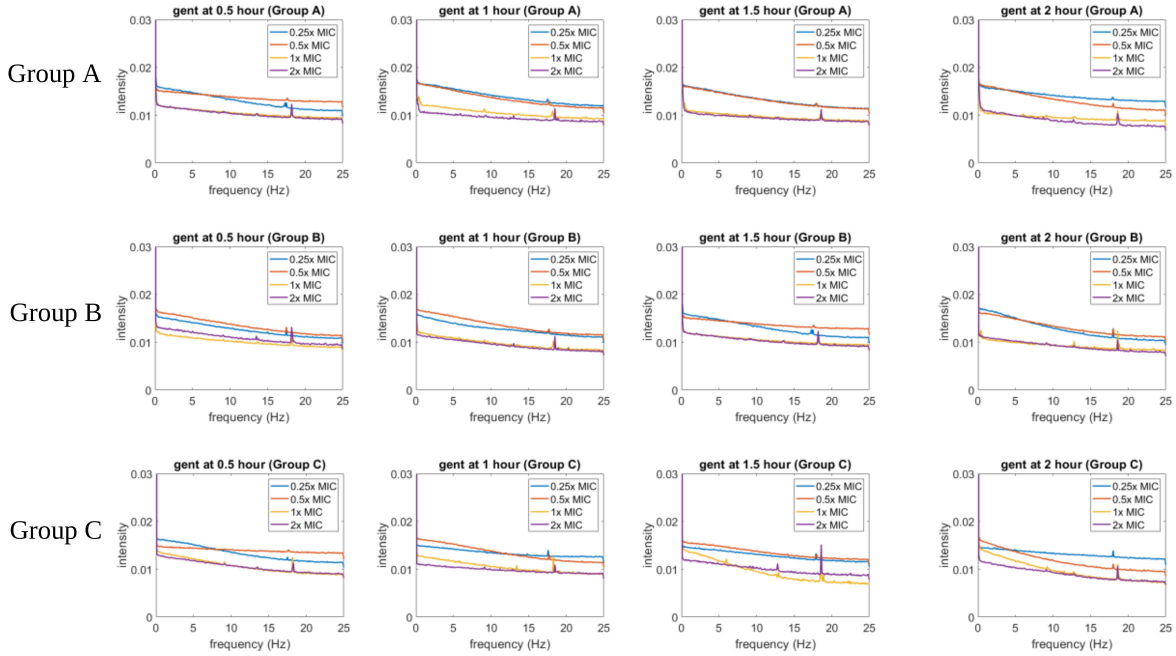


**Fig. S7.** The FT result of the gentamicin group for three independent experiments at time points of 30m, 1h, 90 min, and 2h. The rows correspond to different experiments and the columns indicate different time points.

### Section 4: Confusion matrix for the ANN model and time-evolution

***Table S2.*** *Confusion matrix for ampicillin data at (a) $t_1$, (b) $t_2$ ,(c) $t_3$, (d) $t_4$ at the training process before testing on an independet experiments. The green box is the percentage of the correct pixel-level prediction, while red indicates the incorrect pixel-level prediction. The training has early stopping when the validation error deviate from the training error, the learning algorithm stop and save the parameter values. For $t_1$, the overall accuracy is 77.4%. Compared to the result of testing on the independent experiments, the algorithm can not make accurate prediction. However, at $t_2$, $t_3$, $t_4$, the prediction accuracy is 89%, 95.2%, 95.8% respectively. And we observe correct predtions for independent experiments.*

| Training Confusion Matrix | | | | |
|---|---|---|---|---|
| Output Class | inhibited (0) | 17684 | 8369 | |
| | suceptible (1) | 10432 | 47515 | |
| | | inhibited (0) | suceptible (1) | 77.60% |
| | | Target Class | | 22.40% |

| Validation Confusion Matrix | | | | |
|---|---|---|---|---|
| Output Class | inhibited (0) | 3647 | 1849 | |
| | suceptible (1) | 2268 | 10236 | |
| | | inhibited (0) | suceptible (1) | 77.10% |
| | | Target Class | | 22.90% |

| Test Confusion Matrix | | | | |
|---|---|---|---|---|
| Output Class | inhibited (0) | 3602 | 1831 | |
| | suceptible (1) | 2367 | 10200 | |
| | | inhibited (0) | suceptible (1) | 76.70% |
| | | Target Class | | 22.40% |

| All Confusion Matrix | | | | |
|---|---|---|---|---|
| Output Class | inhibited (0) | 24933 | 12049 | |
| | suceptible (1) | 15067 | 67951 | |
| | | inhibited (0) | suceptible (1) | 77.40% |
| | | Target Class | | 22.60% |

(a)

| Training Confusion Matrix | | | | |
|---|---|---|---|---|
| Output Class | inhibited (0) | 23766 | 5133 | |
| | suceptible (1) | 4108 | 50993 | |
| | | inhibited (0) | suceptible (1) | 89.00% |
| | | Target Class | | 11.00% |

| Validation Confusion Matrix | | | | |
|---|---|---|---|---|
| Output Class | inhibited (0) | 5131 | 1049 | |
| | suceptible (1) | 919 | 10901 | |
| | | inhibited (0) | suceptible (1) | 89.10% |
| | | Target Class | | 10.90% |

| Test Confusion Matrix | | | | |
|---|---|---|---|---|
| Output Class | inhibited (0) | 5152 | 1088 | |
| | suceptible (1) | 924 | 10836 | |
| | | inhibited (0) | suceptible (1) | 88.80% |
| | | Target Class | | 11.20% |

| All Confusion Matrix | | | | |
|---|---|---|---|---|
| Output Class | inhibited (0) | 34049 | 7270 | |
| | suceptible (1) | 5951 | 72730 | |
| | | inhibited (0) | suceptible (1) | 89.00% |
| | | Target Class | | 11.00% |

(b)

| Training Confusion Matrix | | | | |
|---|---|---|---|---|
| Output Class | inhibited (0) | 26359 | 2248 | |
| | suceptible (1) | 1663 | 53730 | |
| | | inhibited (0) | suceptible (1) | 95.30% |
| | | Target Class | | 4.70% |

| Validation Confusion Matrix | | | | |
|---|---|---|---|---|
| Output Class | inhibited (0) | 5620 | 530 | |
| | suceptible (1) | 393 | 11457 | |
| | | inhibited (0) | suceptible (1) | 94.90% |
| | | Target Class | | 5.10% |

| Test Confusion Matrix | | | | |
|---|---|---|---|---|
| Output Class | inhibited (0) | 5618 | 532 | |
| | suceptible (1) | 347 | 11503 | |
| | | inhibited (0) | suceptible (1) | 95.10% |
| | | Target Class | | 4.90% |

| All Confusion Matrix | | | | |
|---|---|---|---|---|
| Output Class | inhibited (0) | 37597 | 3310 | |
| | suceptible (1) | 2403 | 76690 | |
| | | inhibited (0) | suceptible (1) | 95.20% |
| | | Target Class | | 4.80% |

(c)

| Training Confusion Matrix | | | | |
|---|---|---|---|---|
| Output Class | inhibited (0) | 26395 | 2092 | |
| | suceptible (1) | 1443 | 54070 | |
| | | inhibited (0) | suceptible (1) | 95.80% |
| | | Target Class | | 4.20% |

| Validation Confusion Matrix | | | | |
|---|---|---|---|---|
| Output Class | inhibited (0) | 5803 | 461 | |
| | suceptible (1) | 315 | 11421 | |
| | | inhibited (0) | suceptible (1) | 95.70% |
| | | Target Class | | 4.30% |

| Test Confusion Matrix | | | | |
|---|---|---|---|---|
| Output Class | inhibited (0) | 5720 | 456 | |
| | suceptible (1) | 324 | 11500 | |
| | | inhibited (0) | suceptible (1) | 95.70% |
| | | Target Class | | 4.30% |

| All Confusion Matrix | | | | |
|---|---|---|---|---|
| Output Class | inhibited (0) | 37918 | 3009 | |
| | suceptible (1) | 2082 | 76991 | |
| | | inhibited (0) | suceptible (1) | 95.80% |
| | | Target Class | | 4.20% |

(d)

***Table S3.*** *Confusion matrix for gentamicin at (a)* $t_1$*, (b)* $t_2$ *,(c)* $t_3$*, (d)* $t_4$ *at the training process before testing on the independet experiments. The green box is the percentage of the correct pixel-level prediction, while red indicates the incorrect pixel-level prediction. The training has early stopping when the validation error deviates from the training error, and the learning algorithm stop and save the parameter values. The accruay increases from* $t_1$ *of 72.8% to* $t_4$ *of 78.9%.*

| Training Confusion Matrix | | | | |
|---|---|---|---|---|
| Output Class | inhibited (0) | 40518 | 14915 | |
| | suceptible (1) | 15408 | 41159 | |
| | | inhibited (0) | suceptible (1) | 72.90% |
| | | Target Class | | 27.10% |

| Validation Confusion Matrix | | | | |
|---|---|---|---|---|
| Output Class | inhibited (0) | 8694 | 3229 | |
| | suceptible (1) | 3352 | 8725 | |
| | | inhibited (0) | suceptible (1) | 72.60% |
| | | Target Class | | 27.40% |

| Test Confusion Matrix | | | | |
|---|---|---|---|---|
| Output Class | inhibited (0) | 8693 | 3233 | |
| | suceptible (1) | 3335 | 8739 | |
| | | inhibited (0) | suceptible (1) | 72.60% |
| | | Target Class | | 27.40% |

| All Confusion Matrix | | | | |
|---|---|---|---|---|
| Output Class | inhibited (0) | 57905 | 21377 | |
| | suceptible (1) | 22095 | 58623 | |
| | | inhibited (0) | suceptible (1) | 72.80% |
| | | Target Class | | 27.20% |

(a)

| Training Confusion Matrix | | | | |
|---|---|---|---|---|
| Output Class | inhibited (0) | 41411 | 12715 | |
| | suceptible (1) | 14496 | 43378 | |
| | | inhibited (0) | suceptible (1) | 75.70% |
| | | Target Class | | 24.30% |

| Validation Confusion Matrix | | | | |
|---|---|---|---|---|
| Output Class | inhibited (0) | 8853 | 2751 | |
| | suceptible (1) | 3184 | 9212 | |
| | | inhibited (0) | suceptible (1) | 75.30% |
| | | Target Class | | 24.70% |

| Test Confusion Matrix | | | | |
|---|---|---|---|---|
| Output Class | inhibited (0) | 8936 | 2703 | |
| | suceptible (1) | 3120 | 9241 | |
| | | inhibited (0) | suceptible (1) | 75.70% |
| | | Target Class | | 24.30% |

| All Confusion Matrix | | | | |
|---|---|---|---|---|
| Output Class | inhibited (0) | 59200 | 18169 | |
| | suceptible (1) | 20800 | 61831 | |
| | | inhibited (0) | suceptible (1) | 75.60% |
| | | Target Class | | 24.40% |

(b)

| Training Confusion Matrix | | | | |
|---|---|---|---|---|
| Output Class | inhibited (0) | 43097 | 11594 | |
| | suceptible (1) | 12804 | 44505 | |
| | | inhibited (0) | suceptible (1) | 78.20% |
| | | Target Class | | 21.80% |

| Validation Confusion Matrix | | | | |
|---|---|---|---|---|
| Output Class | inhibited (0) | 9206 | 2525 | |
| | suceptible (1) | 2884 | 9385 | |
| | | inhibited (0) | suceptible (1) | 77.50% |
| | | Target Class | | 22.50% |

| Test Confusion Matrix | | | | |
|---|---|---|---|---|
| Output Class | inhibited (0) | 9224 | 2412 | |
| | suceptible (1) | 2785 | 9579 | |
| | | inhibited (0) | suceptible (1) | 78.30% |
| | | Target Class | | 21.70% |

| All Confusion Matrix | | | | |
|---|---|---|---|---|
| Output Class | inhibited (0) | 61527 | 16531 | |
| | suceptible (1) | 18473 | 63469 | |
| | | inhibited (0) | suceptible (1) | 78.10% |
| | | Target Class | | 21.90% |

(c)

| Training Confusion Matrix | | | | |
|---|---|---|---|---|
| Output Class | inhibited (0) | 43362 | 11279 | |
| | suceptible (1) | 12453 | 44906 | |
| | | inhibited (0) | suceptible (1) | 78.80% |
| | | Target Class | | 21.20% |

| Validation Confusion Matrix | | | | |
|---|---|---|---|---|
| Output Class | inhibited (0) | 9470 | 2343 | |
| | suceptible (1) | 2658 | 9529 | |
| | | inhibited (0) | suceptible (1) | 79.20% |
| | | Target Class | | 20.80% |

| Test Confusion Matrix | | | | |
|---|---|---|---|---|
| Output Class | inhibited (0) | 9382 | 2408 | |
| | suceptible (1) | 2675 | 9535 | |
| | | inhibited (0) | suceptible (1) | 78.80% |
| | | Target Class | | 21.20% |

| All Confusion Matrix | | | | |
|---|---|---|---|---|
| Output Class | inhibited (0) | 62214 | 16030 | |
| | suceptible (1) | 17786 | 63970 | |
| | | inhibited (0) | suceptible (1) | 78.90% |
| | | Target Class | | 21.10% |

(d)

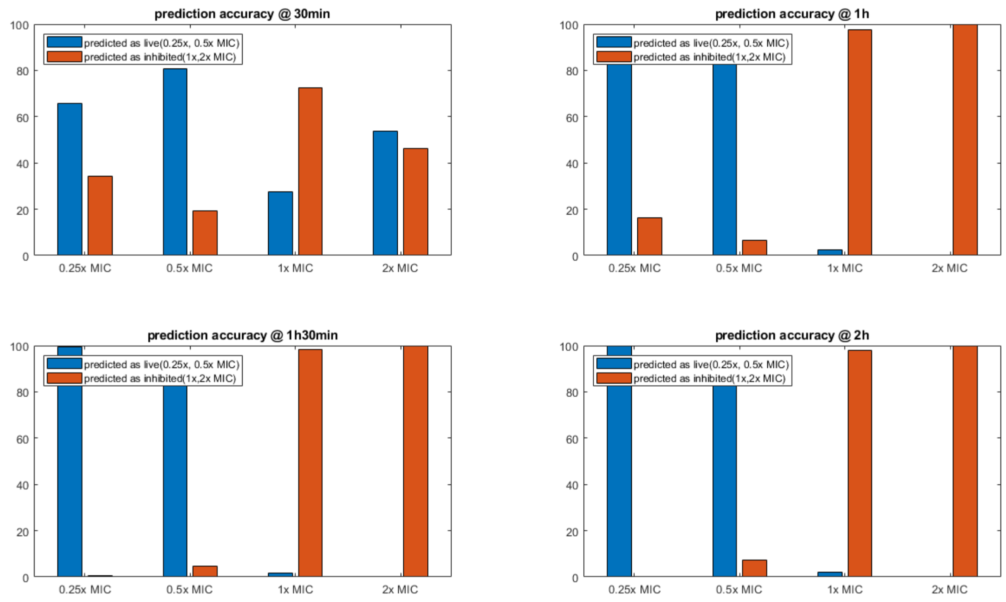


**Fig. S8.** The machine learning prediction results for ampicillin using $t_1$ =30 min data, $t_2$ =60 min, $t_3$ =90 min, and $t_4$ =120 min data. After 60 min, the method can identify MIC with confidence.

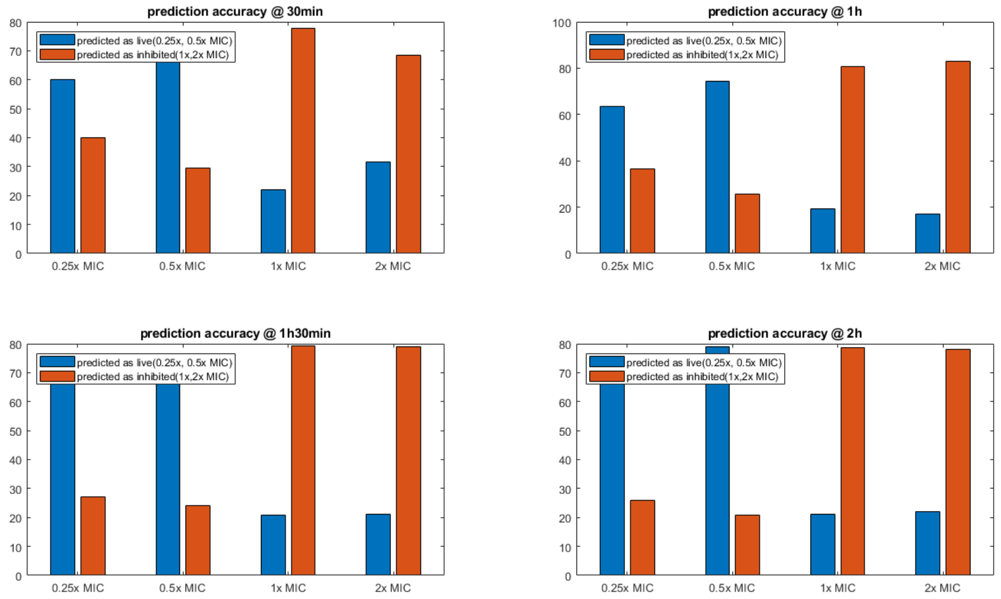


**Fig. S9.** The machine learning prediction results for gentamicin using $t_1$ =30 min data, $t_2$ =60 min, $t_3$ =90 min, and $t_4$ =120 min data. After 60 min, the method can identify MIC with confidence.
